# Automatic calibration of a functional-structural wheat model using an adaptive design and a metamodelling approach

**DOI:** 10.1101/2021.07.29.454328

**Authors:** Emmanuelle Blanc, Jérôme Enjalbert, Pierre Barbillon

## Abstract

- Background and Aims Functional-structural plant models are increasingly being used by plant scientists to address a wide variety of questions. However, the calibration of these complex models is often challenging, mainly because of their high computational cost. In this paper, we applied an automatic method to the calibration of WALTer: a functional-structural wheat model that simulates the plasticity of tillering in response to competition for light.
- Methods We used a Bayesian calibration method to estimate the values of 5 parameters of the WALTer model by fitting the model outputs to tillering dynamics data. The method presented in this paper is based on the Efficient Global Optimisation algorithm. It involves the use of Gaussian process metamodels to generate fast approximations of the model outputs. To account for the uncertainty associated with the metamodels approximations, an adaptive design was used. The efficacy of the method was first assessed using simulated data. The calibration was then applied to experimental data.
- Key Results The method presented here performed well on both simulated and experimental data. In particular, the use of an adaptive design proved to be a very efficient method to improve the quality of the metamodels predictions, especially by reducing the uncertainty in areas of the parameter space that were of interest for the fitting. Moreover, we showed the necessity to have a diversity of field data in order to be able to calibrate the parameters.
- Conclusions The method presented in this paper, based on an adaptive design and Gaussian process metamodels, is an efficient approach for the calibration of WALTer and could be of interest for the calibration of other functional-structural plant models.

## INTRODUCTION

Modelling is a powerful tool to study complex systems. Numerical experiments are often used when real experiments are too difficult or too expensive to carry out. Furthermore, computer models allow to strictly control the conditions of the experiments and to explore conditions beyond what can be covered experimentally. The advantages of computer models have made them essential tools in many disciplines. For example, modelling is widely used in plant sciences and agronomy and has been identified as a promising tool to tackle some of the challenges associated with major issues such as food security (Christensen et al., 2018). In particular, functional-structural plant models (FSPM; Godin and Sinoquet, 2005; Vos et al., 2010) have been more and more used since the 1990s, and they have become a major research subject (Guo et al., 2011; Sievänen et al., 2014; Evers et al., 2018). These individual-based models explicitly represent the plant architecture in 3D and integrate knowledge in ecophysiology and developmental biology. Thus, FSPM take into account the complex interactions between environmental factors, plant development, plant architecture and the underlying physiological processes. These models allow the integration of different scales (from gene to community) to study complex systems in plant science (Louarn and Song, 2020). Thereby, FSPM have been used to study a wide variety of species (both wild and cultivated) and a broad range of questions. Examples of application include prediction of fire behaviour in tree crowns (Parsons et al., 2011); study of the evolutionary emergence of life history strategies along gradients of stress intensity and disturbance frequency (Bornhofen et al., 2011); and study of the impact of plant height on the control of rain-borne diseases in wheat cultivar mixtures (Vidal et al., 2018).

In order to obtain reliable results from FSPM, or from any model, their input parameters must be accurately set. Indeed, the quality of the parameter estimation is critical for the quality of the model predictions. Some parameters are known biological traits that can be directly fixed based on the literature or on experimental data. However, some other parameters cannot be directly measured and are not available in the literature. The values of these parameters have to be indirectly estimated via calibration, which means that the overall model has to be fitted to experimental data. Calibration can be done either manually or automatically. The latter method relies on the use of a search algorithm to identify the “optimal” values of the specified parameters based on the minimization of the distance between the model outputs and the observed data. Automatic calibration has a number of advantages over manual calibration. Indeed, automatic calibration is less subjective than manual calibration, as its success is less dependent on the experience of the modeller (Muleta and Nicklow, 2005). Furthermore, automatic calibration may allow the modeller to look for a distribution of likely values of the parameters instead of looking for an optimal parameter value. Bayesian calibration (Kennedy and O’Hagan, 2001; Higdon et al., 2004; Bayarri et al., 2007) is an automatic calibration method which also provides the modeller with a distribution of values of the parameters that are likely to reproduce the experimental data. The experimental data and the FSPM are linked in a statistical model from which a likelihood is then derived. A prior distribution, which may encode some expert or literature information on the parameter values, is chosen. Since the corresponding posterior distribution is not tractable, MCMC algorithms are run to obtain a sample in the posterior distribution. However, these algorithms resort to many runs of the model, which is why, for complex models, including FSPM, calibration is often done manually (see Lecarpentier et al., 2019 and Gauthier et al., 2020 for example). Indeed, FSPM runs are usually time-consuming and parallel computing of each simulation is complicated because of the interactions between plants in these individual-based models. The solution proposed in the context of Bayesian calibration is to make recourse to a Gaussian Process (Currin et al., 1991; Sacks et al., 1989) metamodel, also known as Kriging metamodel, which provides not only a fast approximation of the FSPM but also a measure of uncertainty concerning the quality of the approximation of the FSPM. Metamodels are widely used to approximate complex models and have sometimes been used with FSPM to allow analyses that would require an important number of simulations (Perez et al., 2018; Da Silva et al., 2014). However, as metamodels are an approximation of the model, the calibration can be less precise and less efficient than with the actual FSPM. To address this problem, a sequential method aiming at improving the Kriging metamodel precision and based on the EGO algorithm (Jones et al., 1998) can be implemented with respect to the calibration goal, as proposed in Damblin et al. (2018). This method has been used for the calibration of complex models (see Carmassi et al., 2019 for example) but has not yet, to the extent of our knowledge, been applied to FSPM. Moreover, the application of this type of method to the FSPM context is particular since it can require to fit several metamodels in parallel to take into account the complex and / or dynamic outputs of the model.

Here, we present how Kriging metamodels and an adaptive design can be used for the calibration of the WALTer FSPM.

## MATERIALS AND METHODS

The efficiency of an automated calibration algorithm to estimate 5 critical parameters of the FSPM WALTer was assessed on both simulated and experimental datasets.

All the analyses were done using the R software (R Core Team, 2017).

### WALTer: a 3D wheat model

WALTer (Lecarpentier et al., 2019) is a functional structural plant model (FSPM) (Vos et al., 2010; Godin & Sinoquet, 2005). This individual-based model simulates the development of the aerial architecture of winter wheat (*Triticum aestivum* L.) from sowing to maturity with a daily time step. In the model, the vegetative development of the plants follows a thermal time schedule and is based on a formalism derived from ADEL-Wheat (Fournier et al., 2003). Thanks to a radiative model (CARIBU: Chelle et al., 1998), WALTer simulates the competition for light between plants and the resulting plasticity of tillering (i.e. the branching ability of grasses). The regulation of tillering in the model is based on three simple rules. (i) Tillers emerge according to empirically fixed probabilities. Then, (ii) an early neighbour perception controls the cessation of tillering: plants stop emitting tillers when the surrounding Green Area Index (GAI: ratio of photosynthetic surface to ground area) reaches a critical value (GAI_c_). Finally, (iii) some of the tillers that were emitted regress: a tiller regresses if the amount of photosynthetically active radiation (PAR) it intercepts falls below a threshold (PAR_t_). During tiller regression, there is a protection period (Δ_prot_) between the deaths of two successive tillers on a plant.

Based on this formalism, WALTer produces useful outputs, such as the tillering dynamics of each plant (i.e. the number of axes on a plant for each day of the simulation). The model has shown its ability to accurately simulate the different patterns of tillering dynamics resulting from variations in sowing density (Lecarpentier et al., 2019).

WALTer is not a completely deterministic model, as several elements in the model integrate some stochasticity. Indeed, the position of the plants, the orientation of the organs, the duration before plant emergence and the emergence of tillers are partly random. Furthermore, the final number of leaves on the main stem 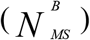 is defined at the scale of the field: it is a decimal number and the decimal part indicates the proportion of plants carrying round 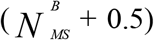 leaves on the main stem. For example, a field with = 11.6 would have 40% of plants with 11 leaves on their main stem and 60% of plants with 12 leaves.

Since its publication in 2019 (Lecarpentier et al.), WALTer has undergone some changes aiming at reducing the computational cost of the simulations, enhancing its realism and improving its ability to simulate mixtures of varieties (Blanc et al., in prep). In particular, the new version of WALTer integrates: a representation of curved leaves (Dornbusch et al., 2009; Fournier and Pradal, 2012; Perez et al., 2016); (ii) an improved discretization of the sky (den Dulk, 1989; Alinea.ASTK, version 2.1.0, 2019); (iii) the possibility to simulate an infinite periodic canopy allowing to discard border effects; (iv) removal of non-visible organs from the 3D representation of plants.

WALTer is available as an open source Python package on the OpenAlea platform (https://github.com/openalea/walter).

### Parameter estimation

Five parameters of the model (Table 1) were estimated by fitting the tillering dynamics. These five parameters were selected based on the results of a global sensitivity analysis of WALTer [**Supplementary Information**] because they had an important impact on the variance of the tillering dynamics and/or because they could not be measured or estimated otherwise.

**Table 1.** List of the 5 WALTer parameters to be estimated, definition, ranges of variation explored for the calibration, values used to generate the simulated test-dataset and units.

| Parameter | Description | Range | Value for the simulated data | Unit |
| --- | --- | --- | --- | --- |
| $GAI_c$ | Green Area Index threshold above which the emission of tillers stops | 0.25 - 1.25 | 0.75 | - |
| $PAR_t$ | PAR threshold below which a tiller does not survive | $10^5$ - $10^6$ | $3 \times 10^5$ | $\mu\text{mol} \cdot \text{cm}^{-2} \cdot ^\circ\text{Cd}^{-1}$ |
| $\Delta_{\text{prot}}$ | Thermal time interval during which two tillers of the same plant cannot die | 10 - 100 | 25 | $^\circ\text{Cd}$ |
| $N_{MS}^B$ | Final number of leaves on the main stem | 8 - 16 | 12.5 | - |
| $L_{\text{max}}^B$ | Final length of the longest blade of the main stem | 8 - 35 | 16.6 | cm |

The parameters that were not estimated here were set according to the bibliography and the manual calibration described in Lecarpentier *et al.*, 2019 for the winter wheat cultivar ‘Maxwell’. For meteorological data, the original sequence of PAR available in Darwinkel (1978) for Lelystad (The Netherlands) and a sequence of daily temperatures obtained by averaging Lelystad data from 2004 to 2014 were used for the simulations.

In order to perform the calibration of WALTer, we assumed a statistical model linking WALTer to the observed data. We denote by *y(d,t)* an observation of the number of axes per m^2^ for density *d* at time *t* and by *f*^(*d,t*)^ (*x*) the evaluation of WALTer for the same density and time with the input parameters set to *x*. Thus, the assumed statistical model is:

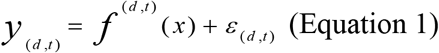

where the distribution of the noise is given by *ε(d,t)~N(0,σ^2^(d,t))* and all the ε are assumed to be independent.

The proposed calibration method is based on the Efficient Global Optimisation (EGO) algorithm (Jones et al., 1998) and involves the following six steps:

1. First, an initial numerical design *D_init_* must be selected to sample the multidimensional parameter space. To ensure a good exploration of the parameter space, a five dimensions maximin Latin Hypercube (LH) design (McKay et al., 1979; Johnson et al., 1990) of 100 parameter-sets was generated with the DiceDesign package (Dupuy et al., 2015) using the ranges detailed in Table 1.
2. The model must then be run over the initial design. All the simulations of the LHD were run with WALTer for *n_d_* = 6 contrasted sowing densities (25, 50, 100, 200, 400 and 800 plants/m^2^), leading to a total of 600 simulations. For each simulation, the mean tillering dynamics of the field was computed by WALTer and a subset of *n*_*t*_ = *13* dates spread across the dynamics was extracted.
3. The initial design must then be used to build a Kriging metamodel which approximates WALTer in the parameter space. For each sowing density and for each of the 13 dates (78 states), a specific metamodel was fitted by using the DiceKriging package (Roustant et al., 2012). For each metamodel, the homogeneous nugget effect was set to WALTer’s variance for the corresponding date and sowing density. WALTer’s variance was computed thanks to 10 replicates of a reference simulation (Table 1). The accuracy of each Kriging metamodel was tested through cross validation by leave-one-out. We denote for a sowing density *d* and a date *t* given the design of numerical experiments D_init_, 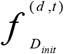 the corresponding Kriging metamodel, the mean of which is denoted by 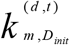 and the variance of which is denoted by 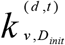.
4. The initial design *D_init_* could then be enriched by the addition of new points. The additional points are selected based on the Expected Improvement (EI) criterion adapted to the calibration goal as in Damblin et al., 2018. This criterion aims to improve the precision of the surrogate model, especially for values of the input parameters which are likely to make a good fit of WALTer to the available experimental data. The difference with Damblin et al., 2018 lies in the fact that several Kriging metamodelds (one for each combination of sowing density and date) are combined. For this step, we decided to enrich the design by the addition of a single point at a time, corresponding to the highest EI criterion among a set of 10 000 points selected from the parameter space by an LH sampling. We denote by *D_k_* the current design of numerical experiments after that *k* points were already added (with the notation *D_0_* = *D_init_*). The EI criterion before adding the *k+1*^th^ point is:

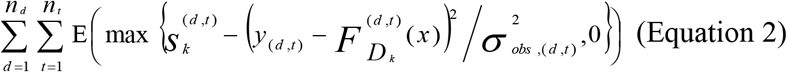

where the expectations are computed with respect to the distributions of the Kriging metamodels 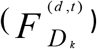 given the current design of experiments (*D_k_*) and 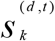 is the current minimal value of the function 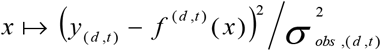 over the current design *D_k_*.
5. A Bayesian calibration method can then be applied to the model. For each input parameter, the posterior distribution was estimated using a Markov Chain Monte Carlo (MCMC) sampling. More precisely, a random walk Metropolis algorithm was used, as implemented in the MCMCpack package (Martin et al., 2011). From Eq.1, the log-likelihood is then computed for any value of the parameters *x* as:

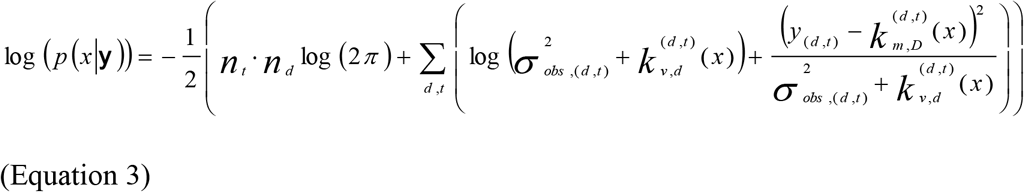

where we denote by 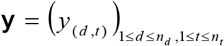 the vector of all observations and *D* the numerical design of experiments that is finally used for performing the Bayesian calibration. The observational variance of each output and the uncertainty of the Kriging metamodel were taken into account to avoid overfitting of the model.
6. The goodness of fit can then be evaluated through cross-validation. For this step, we calibrated the model using only the data from one (200 plants/m^2^) sowing density and used the whole data set for the validation.

### Application to synthetic data

At first, we applied our methodology to a simulated dataset. Using WALTer, we simulated tillering dynamics at six contrasted sowing densities (25, 50, 100, 200, 400 and 800 plants/m^2^) with a fixed set of parameters (Table 1). To build the synthetic dataset, we then extracted 13 dates spread across the tillering dynamics, in accordance with the experimental data described hereafter. Finally, a Gaussian white noise was added to the resulting tillering dynamics. For each date and at each sowing density, the noise had a standard deviation corresponding to WALTer’s standard deviation between replicates at the same date and for the same sowing density. For each sowing density, the observational variance of each date was set to the value of WALTer’s variance for the corresponding date at the same sowing density.

### Application to experimental data

The same methodology was then applied to estimate the five parameter values that would allow the best fitting to experimental data. We used data from an experiment described by Darwinkel (1978), in which the winter wheat cultivar ‘Lely’ was sown in plots of 1 m^2^ at seven contrasted sowing densities (5, 25, 50, 100, 200, 400 and 800 plants/m^2^) in Lelystad (The Netherlands). For each density, the number of shoots per m^2^ was measured at 13 dates spread across the development period of the crop, thus giving a good estimation of the total tillering dynamics.

We decided not to use the data for the lowest sowing density (5 plants/m^2^) because simulations with WALTer for this sowing density would take a very long time. Furthermore, this sowing density represents extreme conditions that would rarely be observed and is thus not the most interesting density to study. Therefore, we focused our calibration on the 6 other sowing densities (25, 50, 100, 200, 400 and 800 plants/m^2^).

For each sowing density, the observational variance, necessary to compute the likelihood (Eq. 1), was set to achieve a coefficient of variation of 20% for the first two dates, of 5% for the last two dates and of 10% for the other dates. These coefficients of variation were chosen to take into account the uncertainty associated with the experimental measurements. A more important coefficient of variation was selected for the first two dates to account for the relatively higher difficulty of measurement at the beginning of the tillering dynamics and for a possible temporal shift due to the discrepancy between the approximated temperature sequence used for the simulations and the real experimental one, not available. The coefficient of variation for the last two dates of the tillering dynamics was set to a smaller value to account for the relative simplicity of measurement on mature plants and for the importance of the fit for these dates as they represent an important component of the yield (number of spikes/m^2^).

## RESULTS

### Application to synthetic data

The accuracy of the calibration is low when it is performed using the Kriging metamodel constructed from the initial design of 100 simulations (Fig 2, top row). Indeed, the posterior distributions of Δ_prot_ and 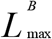 do not include the actual values used to generate the synthetic data. The posterior density for PAR_t_ has a small variance but is shifted from the actual parameter value and the posterior densities of GAI_c_ and 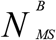 include the actual parameter values but have rather important variances. On the other hand, the accuracy of the calibration is greatly improved by the use of the sequential design, even with an addition of only 15 points. Using this enriched design, the posterior distributions of all input parameters have small variances and include the values used to generate the synthetic data (Fig 2, bottom row). Thus, as shown in Figure 3, the enriched design allows to select a set of parameters that reproduces accurately the simulated tillering dynamics for all sowing densities.

**Fig. 1.**
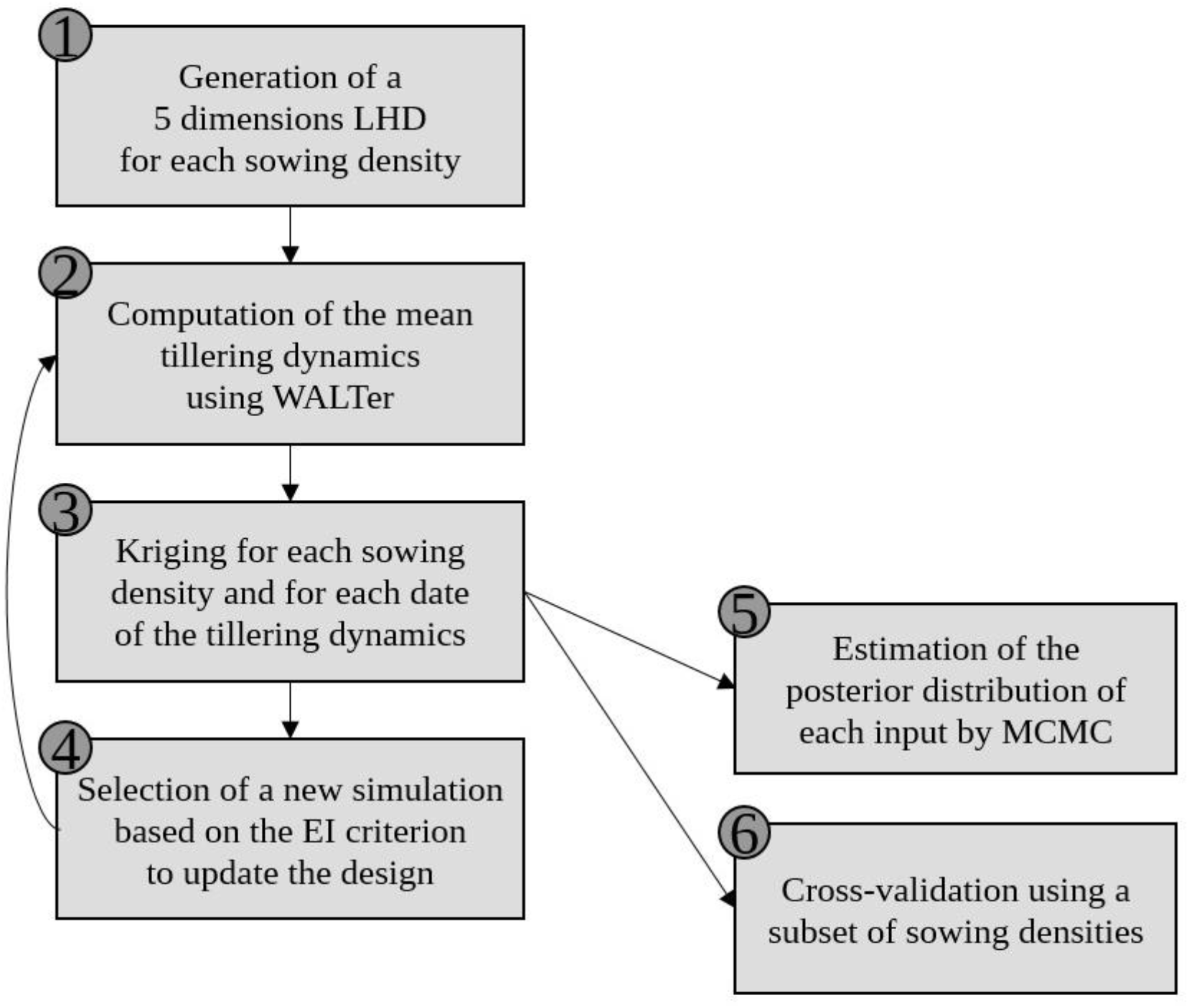
Diagrammatic representation of the proposed calibration method

**Fig. 2.**
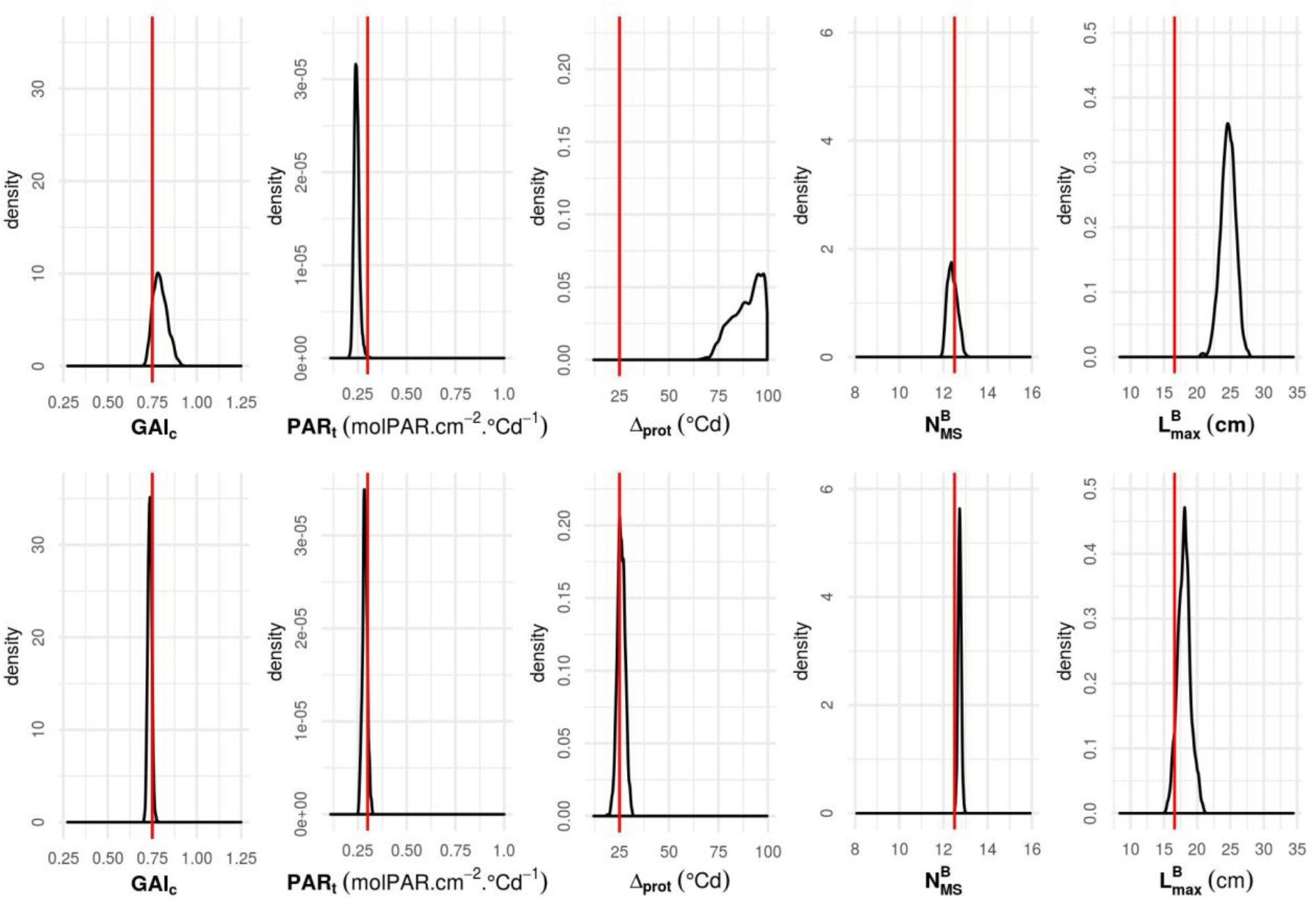
Calibration results using the complete simulated dataset: posterior distributions of WALTer parameters (GAI_c_, PAR_t_, Δ_prot_, 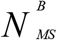 and 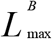) using the simulated data from all densities and the initial LH design of 100 simulations (top row) or the enriched design after 15 loops of EGO (bottom row). Parameter values used to simulate the data are shown with a red vertical line.

**Fig 3.**
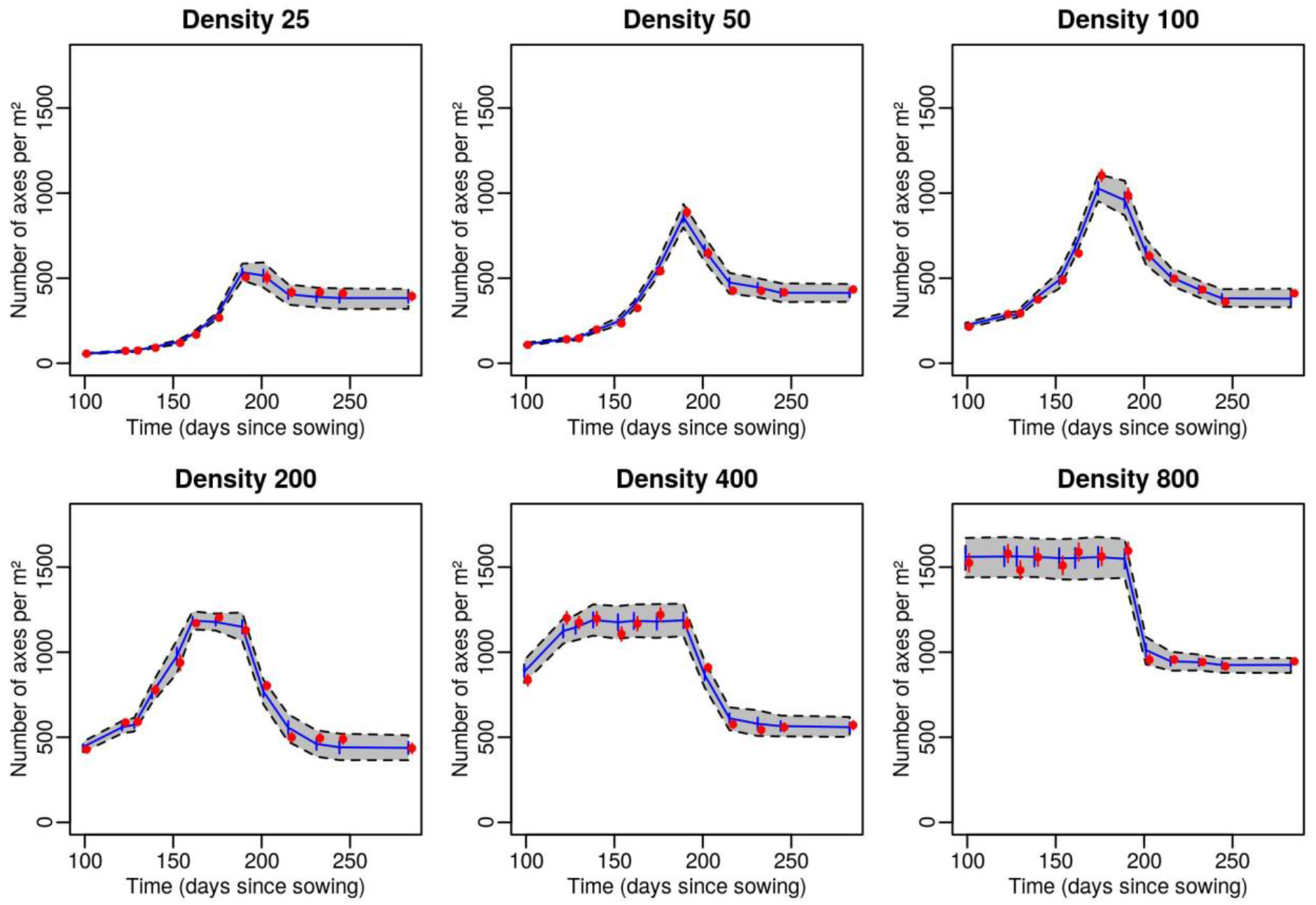
Validation of the calibration using the complete simulated dataset: number of axes per m^2^ vs. time since sowing at six sowing densities (25, 50, 100, 200, 400 and 800 plants.m^−2^). Red dots represent the synthetic data and red segments are the associated observational variance; blue lines represent the mean Kriging prediction for the set of input parameters with the best loglikelihood and blue segments are the associated Kriging standard deviation. The grey area delimited by dotted lines is the 95% credibility interval taking into account the Kriging standard deviation and uncertainty on the calibration parameter. Red points and segments are shifted on the x-axis to avoid overlapping. The set with the best loglikelihood was selected by MCMC using the enriched design after 15 loops of EGO and simulated data from all sowing densities.

However, even when using the design enriched with 15 simulations, the accuracy of the calibration is greatly deteriorated when it is carried out only with the data of a single sowing density (Fig 4). Indeed, the posterior distributions obtained using only the data at 200 plants/m^2^ all include the actual parameter values but their variances are very large. This resulted in some of the simulated data (especially the end of the tillering dynamics at densities 50 and 100 plants/m^2^) being outside the 95% credibility range of the Kriging metamodel when using this single density to select the ‘optimal’ set of parameters (Fig 5). However, the fit to the synthetic data still seems reasonably good, as the 95% credibility range includes the vast majority of the data.

**Fig. 4.**
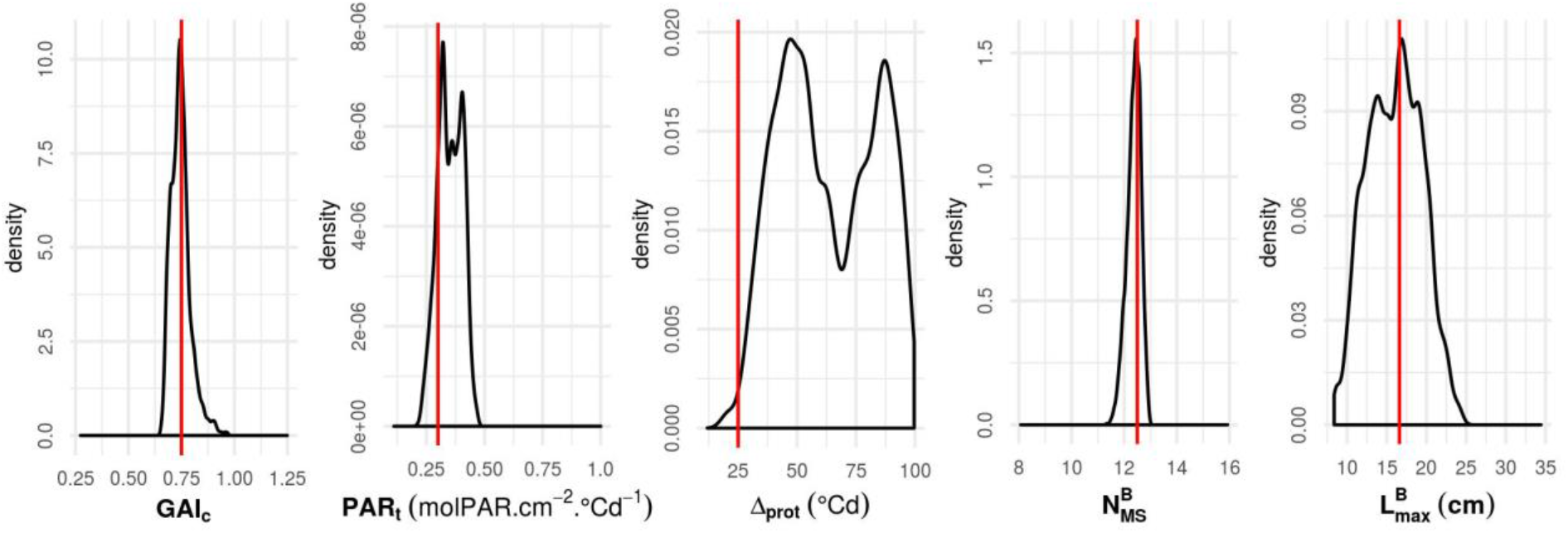
Calibration results using a subset of the simulated dataset: posterior distributions of WALTer parameters (GAI_c_, PAR_t_, Δ_prot_, 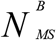 and 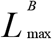) using the enriched design after 15 loops of EGO and simulated data from density 200 plants/m^2^. Parameter values used to simulate the data are shown with a red vertical line.

**Fig 5.**
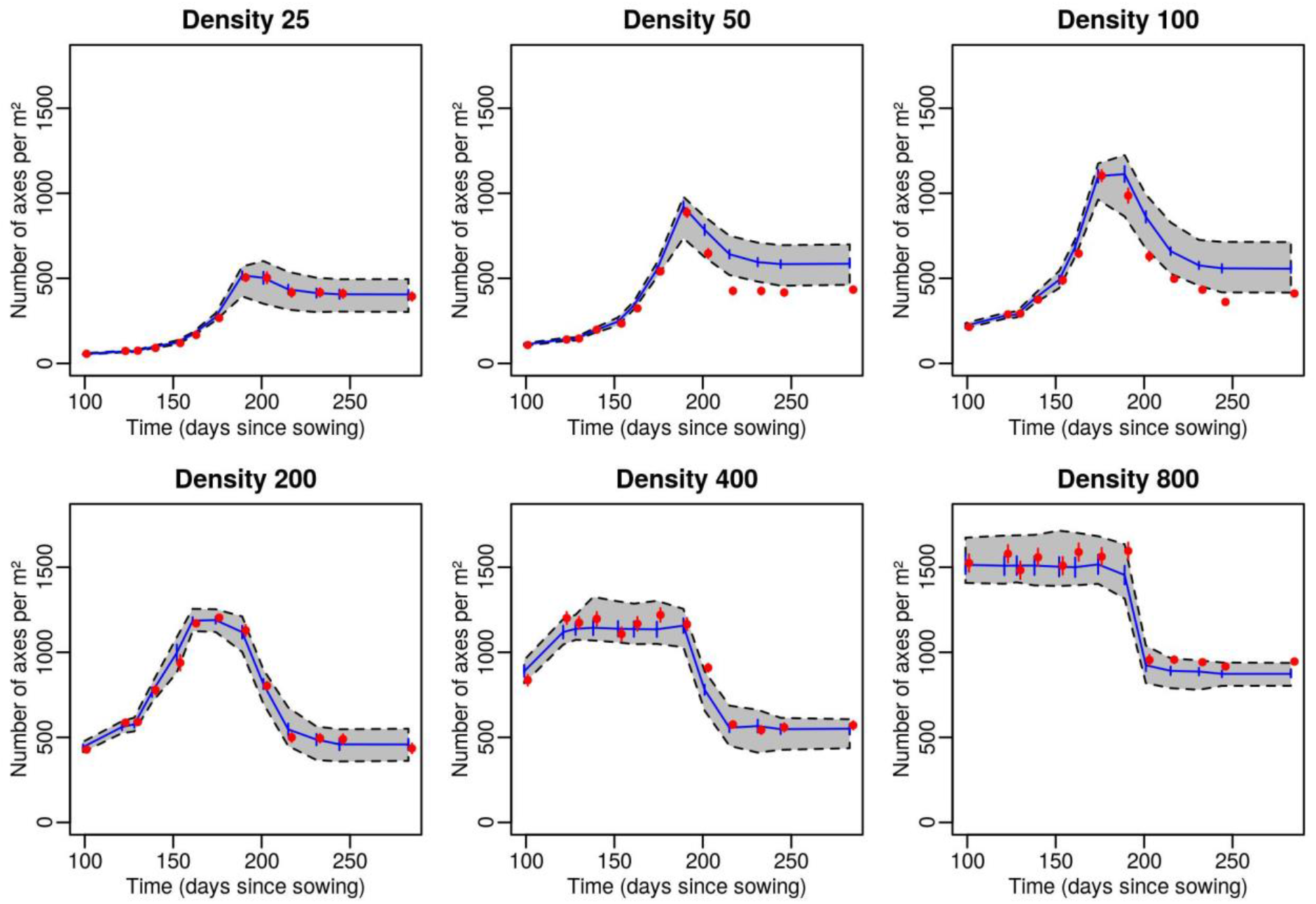
Validation of the calibration using a subset of the simulated dataset: number of axes per m^2^ vs. time since sowing at six sowing densities (25, 50, 100, 200, 400 and 800 plants.m^−2^). Red dots represent the synthetic data and red segments are the associated observational variance; blue lines represent the mean Kriging prediction for the set of input parameters with the best loglikelihood and blue segments are the associated Kriging standard deviation. The grey area delimited by dotted lines is the 95% credibility interval taking into account the Kriging standard deviation and uncertainty on the calibration parameter. Red points and segments are shifted on the x-axis to avoid overlapping. The set with the best loglikelihood was selected by MCMC using the enriched design after 15 loops of EGO and simulated data from density 200 plants/m^2^.

### Application to experimental data

When the enriched design is used to calibrate WALTer on the experimental data from Darwinkel (1978), the posterior distributions (Fig 6) show contrasting results depending on the input parameter considered. The posterior density for 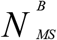 and GAI_c_, which are the 2 parameters with the most influence on the tillering dynamics according to the global sensitivity analysis [**Supplementary Information**], only include intermediate values of the parameters and have a rather low variance. On the other hand, the posterior density for Δ_prot_ has a very large variance and includes almost all the range of values explored. As for the posterior density of PAR_t_, its variance is low, but it only includes values that are close to the lower bound of the variation range. Similarly, the posterior density of 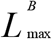 is clearly shifted towards the upper bound of the variation range, even though its variance is quite large.

**Fig. 6.**
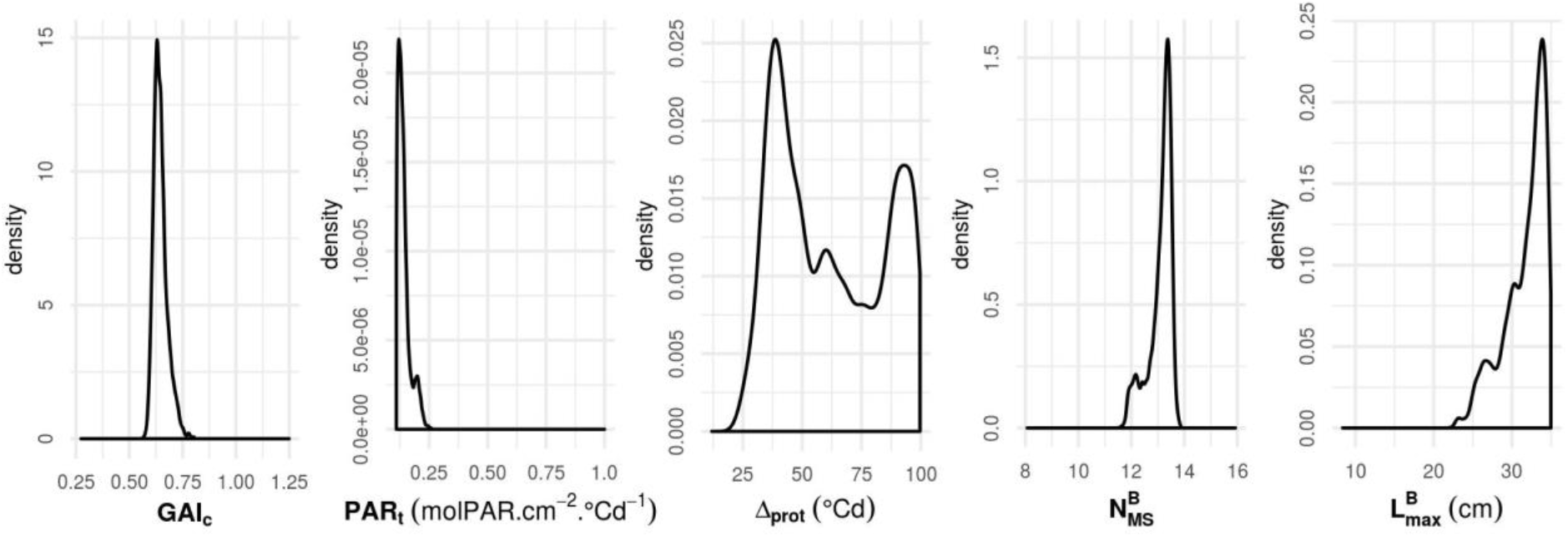
Calibration results using the experimental dataset: posterior distributions of WALTer parameters (GAI_c_, PAR_t_, Δ_prot_, 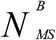 and 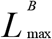) using the experimental data from Darwinkel (1978) at all densities and the enriched design after 15 loops of EGO.

With the ‘optimal’ set of parameters selected by MCMC, there is a rather good fit between the simulated tillering dynamics and the experimental dynamics of Darwinkel (1978) (Fig 7). However, some of the experimental data is outside the 95% credibility range of the Kriging metamodel, even when the observational variance is considered. In particular, for all sowing densities except 800 plants/m^2^, the model fails to reproduce the number of axes per m^2^ observed experimentally for the first 2 dates of the dynamics.

**Fig 7.**
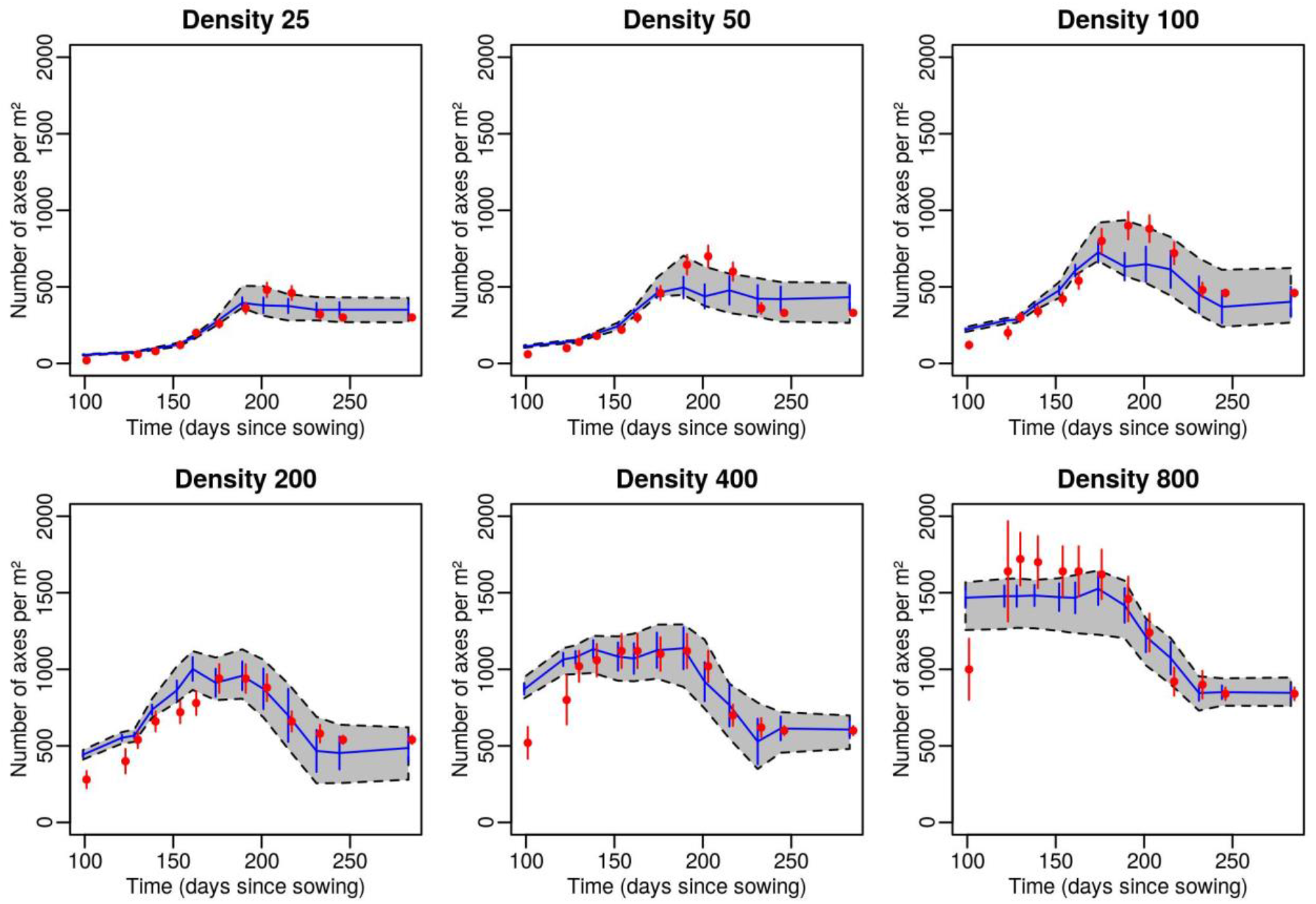
Validation of the calibration using the experimental dataset: number of axes per m^2^ vs. time since sowing at six sowing densities (25, 50, 100, 200, 400 and 800 plants.m^−2^). Red dots represent the experimental data from Darwinkel (1978) and red segments are the associated observational variance; blue lines represent the mean Kriging prediction for the set of input parameters with the best loglikelihood and blue segments are the associated Kriging standard deviation. The grey area delimited by dotted lines is the 95% credibility interval taking into account the Kriging standard deviation. Red points and segments are shifted on the x-axis to avoid overlapping. The set with the best loglikelihood was selected by MCMC using the enriched design after 15 loops of EGO and experimental data of Darwinkel (1978) from all sowing densities.

## DISCUSSION

The calibration of complex models, such as FSPM, involves several challenges, due in part to the high computational cost of the simulations, as well as the numerous parameters to consider. This study illustrates the interest of using a sequential design with a metamodelling approach to calibrate the WALTer FSPM for 5 critical parameters, based on data from the tillering dynamics. The method presented here performed well, especially when applied to synthetic data. The Kriging metamodel efficiently approximated the mean number of axes per m^2^ simulated by WALTer for 13 dates in pure stands. This highlights the interest of using a Kriging metamodel with WALTer to limit the computational cost and thus allow to use methods that require a large number of runs, such as the MCMC. Importantly, a set of 100 simulations was not sufficient to ensure a quality of the metamodel that was high enough for the calibration. However, an enriched design with only 15 additional simulations (115 simulations total), obtained by the sequential method, allowed for a satisfactory fit of the metamodel outputs to both synthetic and experimental data. The use of an adaptive design thus proved to be a very efficient method to improve the quality of the metamodel, reducing the uncertainty in areas of the parameter space that are of interest for the fitting. Contrary to the manual calibration previously applied to WALTer (Lecarpentier et al., 2019), the method presented here provides the practitioner with a distribution of likely values for each parameter considered. Moreover, the Bayesian method presented allows for a more rigorous exploration of the parameter space than the previous manual calibration. However, the method presented here could be improved. For example, the choice of a maximin Latin Hypercube Sampling (LHS) as the initial design for the sequential method can be discussed. Indeed, Zhang et al. (2019) argue that other designs outperform maximin LHS in both static and sequential settings. Furthermore, the good performance of the method for the calibration on the experimental data of Darwinkel (1978) relies on assumptions regarding the observational variance. Indeed, the experimental dataset consisted only of the mean tillering dynamics of the plots and no variance was provided. The uncertainty associated with the experimental measurements was thus assumed to be rather high, but a different observational variance may have impacted the results of the fitting. It is also important to mention the potential divergence between the sequence of daily temperature used in this work and the real one, not available, that has an impact on the quality of the fitting and on its robustness. This highlights the importance of the quality of the data used for the fitting. For FSPM, parameter estimation often requires a lot of data at various scales and the issue of the limited availability of such appropriate datasets is a concern (Louarn and Song, 2020). Interestingly, our study also provides information regarding the type of data necessary for the calibration of WALTer. First of all, our results suggest that data on the tillering dynamics are sufficient to estimate the value of parameters controlling the regulation of tillering (GAI_c_, PAR_t_ and Δ_prot_), but also to estimate the final number of leaves on the main stem 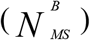 and the final length of the longest blade 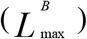. However, it is noteworthy that the two architectural parameters (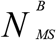 and 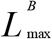) can be measured experimentally. It might thus be possible to estimate the values of the parameters controlling the tillering dynamics (GAI_c_, PAR_t_ and Δ_prot_) with a dataset containing fewer measurement dates, provided that 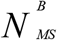 and/or 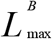 are measured experimentally. Moreover, the value of the five parameters were estimated with a set of 13 dates spread across the development period of the crop. However, since the parameters estimated here do not impact the rate of emission of the tillers, it is possible that the calibration could be done with a slightly lower measurement effort by reducing the number of dates at the beginning of the dynamics. In line with these last considerations, a second interest of the proposed approach is to allow the exploration of alternative experimental schemes at a low computational cost, thanks to the use of metamodels. Our study has for example illustrated the importance of collecting data for several sowing densities for the calibration of WALTer, as the estimation of the parameters is deteriorated when it is based only on data collected at a 200 plants/m^2^ sowing density. Thanks to our simulations, it would thus be possible to identify more precisely the sowing densities that should be used to generate experimental data for the calibration of WALTer. In particular, it would be possible to identify which parameters require data at low or high sowing densities. In future work, we will explore these possibilities based on the synthetic data generated here, to identify the minimal dataset necessary to achieve a satisfactory calibration.

The method presented here, based on an adaptive design and a Kriging metamodel, is an efficient approach for the calibration of WALTer and could be of interest for the calibration of other FSPM. Interestingly, by reducing the computational cost of parameter space exploration, this approach would make it possible to both calibrate FSPM for a large number of genotypes or conditions, and help design the experiments needed to collect the necessary data for a reliable parameter estimation.

## SUPPLEMENTARY INFORMATION

Supplementary information consist of the following. SI1_WALTer_sensitivity_analysis: information regarding the sensitivity analysis of WALTer.

## ACKNOWLEDGEMENTS

The authors thank Christian Fournier, Christophe Lecarpentier and Christophe Pradal for their valuable help in the development of the new version of WALTer. This work is supported by public grants overseen by the French National research Agency (ANR) as part of the “Investissement d’Avenir” program, through the “IDI 2017” project funded by the IDEX Paris-Saclay, ANR-11-IDEX-0003-02 and the “MoBiDiv” project, ANR-20-PCPA-0006.

