## Supplementary information 1 for "Automatic calibration of a functional-structural wheat model using an adaptive design and a metamodelling approach"

**Description of the sensitivity analysis of WALTer**

***Method***

A global sensitivity analysis of WALTer was conducted to evaluate the impact of 13 input parameters and variables of the model (Table SI1) on its outputs. Given the high computational cost of simulations, a fractional factorial design was selected. Three levels were considered for each input factor (Table SI1) and a design of resolution V, composed of 729 simulations, was generated using the R package *planor*. The analysis was done using simulations of 56 plants and each simulation was replicated 5 times to account for the impact of the stochasticity of the model. We denote by *rep*, the random seed used to compute the simulation. For each simulation, the mean tillering dynamics of the plot was extracted from the outputs of WALTer. Thanks to the R package *multisensi*, total sensitivity indices were computed for each date of the tillering dynamics and represented graphically.

Table SI1. Input factors of the sensitivity analysis, description, selected values for the fractional factorial design and units.

| **Input factor** | **Description** | **Values** | | | **Unit** |
|---|---|---|---|---|---|
| Density | Sowing density | 100 | 200 | 400 | plant/m² |
| $GAI_c$ | Green Area Index threshold above which the emission of tillers stops | 0.25 | 0.75 | 1.25 | - |
| $d_{GAIp}$ | Maximal range for plant detection | 0.2 | 0.5 | 1 | m |
| $PAR_t$ | PAR threshold below which a tiller does not survive | $2x10^5$ | $3x10^5$ | $4x10^5$ | $\mu mol.cm^{-2}.°Cd^{-1}$ |
| $\Delta_{prot}$ | Thermal time interval during which two tillers of the same plant cannot die | 10 | 50 | 100 | °Cd |
| $t_{beg}^{reg}$ | Physiological stage at which tiller regression can start | 3.2 | 4.2 | 5.2 | - |
| $N_{MS}^{B}$ | Final number of leaves on the main stem | 8.3 | 11.3 | 15.3 | - |
| $L_{\max}^{B}$ | Final length of the longest blade of the main stem | 8 | 16.6 | 35 | cm |
| Leaf type | Leaf inclination: combination of a parameter for blade insertion angle and a parameter for blade curvature | planophile | medium | erectophile | - |
| $P_T$ | Probability of tiller emergence (except coleoptile tiller) | 0.5 | 0.75 | 0.95 | - |
| $P_{CT}$ | Probability of emergence of the coleoptile tiller | 0 | 0.35 | 0.7 | - |
| Height | Final height of the main stem | 40 | 100 | 160 | cm |
| Phl | Phylochron: thermal time interval between the emergence of two successive leaves | 80 | 100 | 125 | °Cd |

***Results***

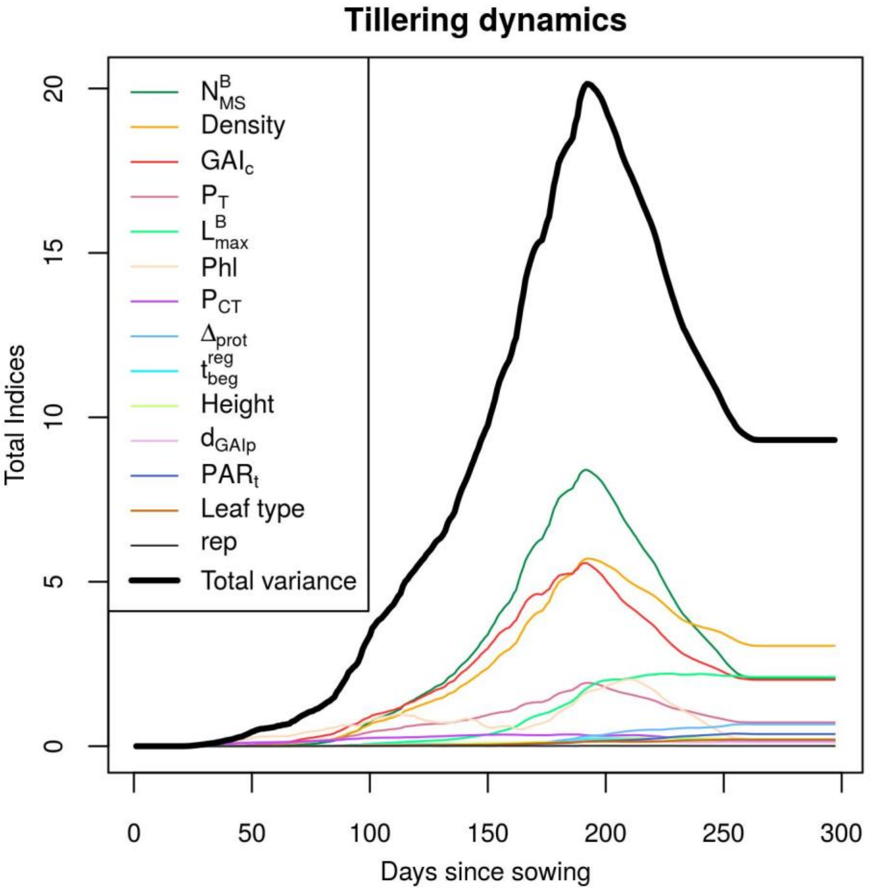


Fig SI1. Total sensitivity indices computed for each date of the tillering dynamics.
